# Regulated development of cannibalistic supergiant cells

**DOI:** 10.1101/2025.08.19.671124

**Authors:** Ben T. Larson, Daniele Giannotti, Mahara Mtawali, Samuel J. Lord, Vittorio Boscaro, Patrick J. Keeling

## Abstract

Virtually all paradigms in developmental biology apply to differentiating cells and tissues within multicellular animals and plants. However, unicellular eukaryotes, which must simultaneously perform all roles necessary for organismal function, also form complex and specialized structures, using processes that take place exclusively within the confines of a single cell. Here, we describe a ciliate (*Euplotes gigatrox* sp. nov.) undergoing drastic morphological transformations within a genetically uniform population, the most spectacular being the appearance of “supergiants” that increase in size, change shape, and modify their locomotion and feeding behaviour to cannibalize clonal relatives. We explore supergiant formation from the perspective of life cycle, ecological strategy, and gene expression, demonstrating that supergiants are distinct, regulated, transcriptionally unique stages. These reversibly differentiated cells require both external and internal triggers to develop and have evolved regulatory loops to ensure coupling between environmental and physiological conditions. This system provides a blueprint for approaching both cell differentiation and functional ecology in unicellular organisms, which might open new avenues for the generalization and contextualization of known morphogenetic mechanisms, as well as the discovery of new ones.

## Introduction

Our understanding of cell development and morphogenesis focuses primarily on the differentiation of cell types within multicellular organisms, and as a result the dominant models for the determination of cell shape and function mostly apply to animal and plant tissues. But the vast majority of eukaryotic diversity resides in unicellular protists^1^, where the cell is the whole organism, facing many functional challenges within the confines of a single plasma membrane: moving, hunting, ingesting prey, escaping predators, finding a mate, but also replicating DNA, remodelling the cytoskeleton, harvesting energy, and managing vesicle trafficking, among other physiological processes^2,3^. The tissues of “complex” multicellular organisms compartmentalize functions and arise from the integration of highly specialized cell types, whose activities are generally each restricted to one primary role^4^. This has become the paradigm for developmental biology, but it is in fact an exception both among extant organisms and throughout the history of life: microbial eukaryotes (protists) must solve most of the same problems with organelles, membranes, and vesicles, rather than organs, tissues, and limbs^5^. In contrast to prokaryotes, which excel at finding metabolic solutions, unicellular eukaryotes rely more on morphological and behavioural features, the complexity of which rivals those of small invertebrates^2,3^. Heterotrophic protists in particular respond to challenges with similar strategies, but completely different tools, than the more familiar animal models, allowing for comparisons between systems replete with both similarities and differences.

Among heterotrophic protists, ciliates are especially useful models because of their size, morphological complexity, and relative ease of cultivation, not to mention the uniqueness, among unicellular organisms, of their germline/soma separation^6^; some have in fact provided insights in fields as disparate as cellular aging, regeneration, biophysics, and epigenetics^7–11^. Ciliates of the genus *Euplotes* have attracted attention since the earliest days of microscopy, due to their ubiquity and striking features. *Euplotes* species occur in most aquatic ecosystems, and their movement, mating habits, symbiotic relationships, biogeography, and adaptations to local environments have all been extensively investigated^12–15^. *Euplotes* cells have a highly ordered, complex animal-like body plan, with cilia packed into larger structures, called membranelles and cirri, that have been modified for feeding (by generating water currents), swimming, or to be used as “legs” for crawling or walking across substrates (Fig. 1A). The morphogenesis of these structures must be faithfully executed at each cell division, all while the cell largely remains active (moving, eating, etc.). This was once the subject of fascinating research that has largely been abandoned in the last 40 years, until recently^16–19^.

**Fig 1.**
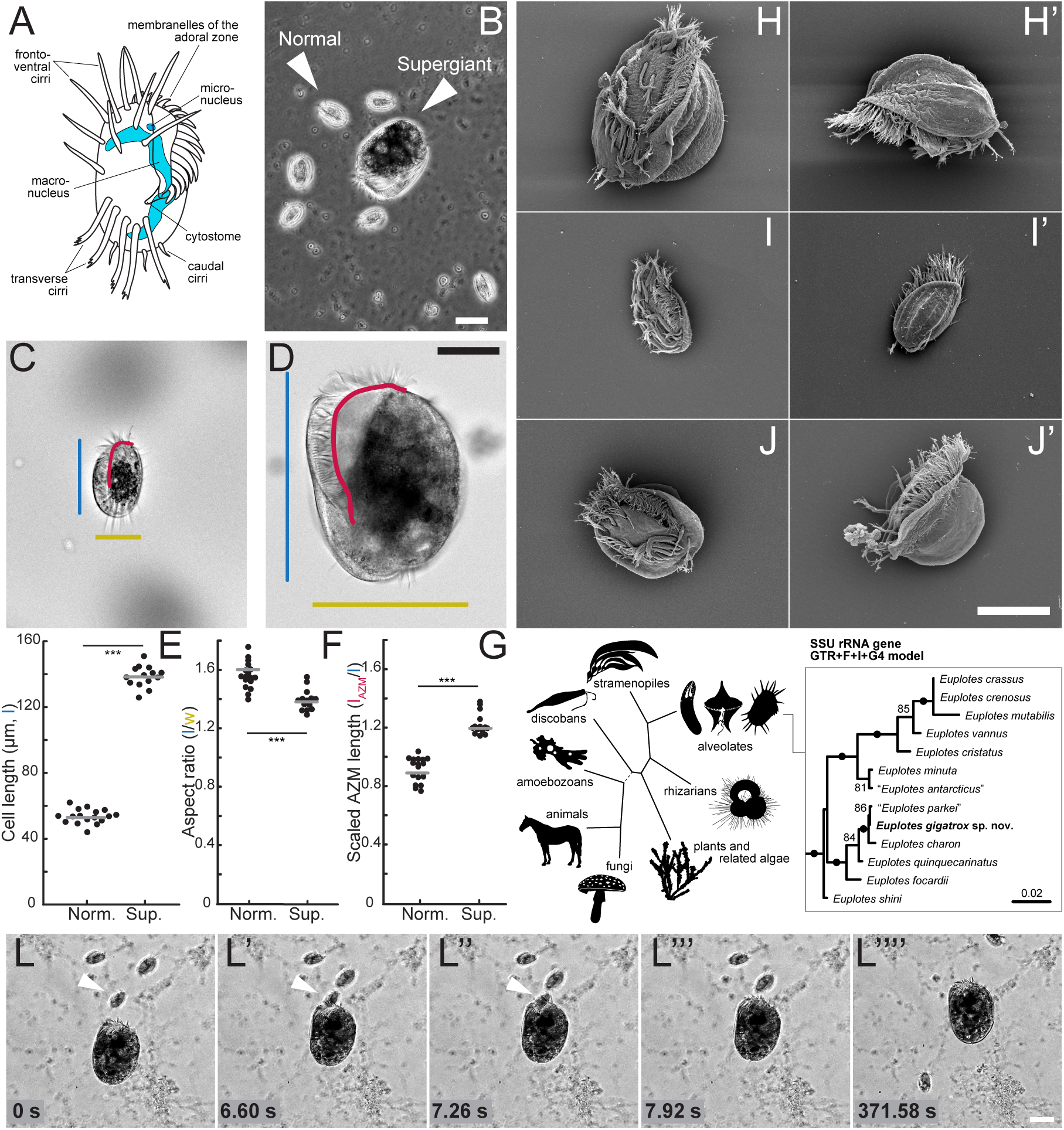
*Euplotes gigatrox* displays marked phenotypic polymorphism including supergiant cannibal cells. (A) A cartoon diagram illustrating basic cellular morphological characteristics of *Euplotes* species, including ventral cirri used for walking locomotion, membranelles (membranelles of the adoral zone, AZM) used to generate flows for swimming and feeding, sensory caudal cirri, and macro- and micronuclei. (B) A phase contrast image displaying genetically identical “normal” and “supergiant” morphs of *E. gigatrox* in close association in a culture flask. Scale bar = 50 µm. (C) Illustration of the morphological characterization of a normal morph with length in blue, width in yellow, and AZM length in red. (D) Illustration of the morphological characterization of a supergiant morph with length in blue, width in yellow, and AZM length in red. Scale bar for both C and D = 50 µm. (E) Comparison of cell lengths between normal morphs and supergiants demonstrates that supergiants are significantly larger (138.2 +/- 4.3 µm versus 53.6 +/- 4.3 µm, mean +/- standard deviation). (F) Comparison of cell aspect ratios (cell length divided by width) between normal morphs and supergiants demonstrates that supergiants have a significantly less elongated cell body (1.4 +/- 0.1 versus 1.6 +/- 0.1, mean +/- standard deviation). (G) Comparison of scaled AZM length (AZM length divided by cell length) between normal morphs and supergiants demonstrates that supergiants have an extended AZM relative to cell size (1.22 +/- 0.08 versus 0.91 +/- 0.09, mean +/- standard deviation). In E-G, measurements are from 17 normal morphs and 14 supergiants, *** indicates *p*<0.001 by two-sample t-test, and gray bars indicate means. (H, H’) Scanning electron micrographs displaying ultrastructural morphology of supergiants. (I, I’) Scanning electron micrographs of normal morphs. (J, J’) Scanning electron micrographs of winged morphs. Scale bar for H-J’ = 50 µm. (K) Phylogenetic placement of *E. gigatrox* in the tree of *Euplotes* (extracted from Fig. S2; numbers close to nodes represent bootstrap support if above 75%, with dots indicating full support), and of *Euplotes* in the tree of eukaryotes. (L-L’’’’) Timelapse images from a video capturing a supergiant predation event on a normal morph prey cell, indicated by the white arrowhead. Scale bar = 50 µm.

One previously reported but seldom explored feature of certain ciliate species, including a few *Euplotes,* is the reversible transformation of cells within a genetically identical population into much larger “supergiants” that cannibalistically consume their relatives^20–24^. The formation of supergiants raises several evolutionary, ecological, ethological, and physiological questions. Here we combine classical observations of morphological, ultrastructural, and behavioural changes with molecular and single-cell transcriptomic methods to show that *Euplotes* supergiants constitute a well-regulated developmental stage: they are differentiated cell types with specific morphology, behaviour, and gene expression profiles, induced both by external stimuli, as previously speculated^22,23,25^, but also by their internal state. These features make *Euplotes* supergiants unique models for the comparative study of developmental biology mechanisms in unicellular organisms.

## Results and Discussion

### *Euplotes gigatrox* sp. nov. displays an extremely polymorphic phenotype

While surveying protist diversity on the Caribbean island of Curaçao, we collected *Euplotes* cells from a seawater filtration system that draws its supply from near-shore waters. The ciliates were initially co-isolated with other microbial eukaryotes and bacteria. To establish laboratory cultures, we manually picked *Euplotes* cells and placed them in nutrient-supplemented artificial seawater along with co-occurring bacteria, which serve as a food source. Cultures were kept at 20 °C and passaged approximately once per month. After several months, especially large cells began to appear sporadically (supergiants, Fig. 1B). These supergiants are morphologically distinguishable from the abundant “normal” cells by their large size (cell length = 138.2 +/- 6.4 µm versus 53.6 +/- 4.3 µm), less elongated cell bodies (cell aspect ratio = 1.4 +/- 0.1 versus 1.6 +/- 0.1), and extended adoral zone of membranelles (scaled AZM length = 1.22 +/- 0.08 versus 0.91 +/- 0.09, Fig. 1C-I). Despite differential scaling of cell size and shape, we observed that supergiants retained the same number of ventral cirri (bundles of cilia) as normal morphs (Fig. 1H-I’). Upon closer inspection, we also noted further morphological diversity in the cultures, including “winged” morphs (Fig. 1J), which might serve a defensive purpose^26,27^, as well as cells that appeared intermediate between the normal and supergiant phenotype (Fig. S1). It was impossible to unambiguously categorize intermediate cells into well-defined, cohesive groups based on gross morphology, and we hypothesized that they represent progressive stages in supergiant development.

Phylogenetic analysis of the small subunit (SSU) ribosomal RNA gene and morphological comparisons confirmed that this *Euplotes* isolate belonged to a previously undescribed species (Fig. 1K; see Text S1, Fig. S2 for a more detailed species characterization), which we name *Euplotes gigatrox,* in reference to the presence of fierce, supergiant cells.

### “Supergiants” exhibit altered behavioural patterns

Giant cells have been observed in many ciliates, and are often associated with cannibalistic feeding^22,24,25^. The supergiants of *E. gigatrox* were also observed to consume clonal relatives (Fig. 1L, Videos S1-S2) at a rate of up to approximately 1 prey every 10 min. In cannibalistic feeding, predator cells “run over” normal morphs until they are lodged in the oral cavity, where they are engulfed. This contrasts sharply with filter feeding in normal morphs and other *Euplotes* species, where a current is generated by the membranelles to pull bacteria and small protists in. The behavioural and structural differences between normal cells and supergiant cannibals offer fascinating eco-evolutionary and developmental implications, so we sought to characterize them in more detail.

We found that supergiants exhibit distinct motility patterns. Freely behaving normal morphs under dilute environmental conditions display the expected range of cell movements^15,28^, including walking along the substrate at various speeds as well as swimming along helical trajectories in fluid (Fig. 2A, Video S3). Supergiants, in contrast, only walk, often along curling trajectories, and spontaneous swimming was never observed (Fig. 2B,C, Video S4). Instantaneous velocities of walking supergiants show generally higher speeds, but also a much broader distribution compared to the speeds of normal morphs, despite the fact that the latter can both walk and rapidly swim (see Methods, Fig. 2C). The turning angle distribution (see Methods) of supergiants displays a shoulder at around 30 degrees due to the persistent curvature of many trajectories, while the turning angle distribution for normal morphs exhibits a single peak and no shoulder, consistent with prior analyses of other *Euplotes* species^15^ (Fig. 2D). We also examined the mean square displacement (MSD) of cell movement patterns (see Methods), which is commonly used to characterize trajectories^29^. Normalized MSD curves for both morphs display similar initial scaling exponents of 1.43 (95% confidence interval bounds: 1.41 - 1.46; R^2^ = 0.9977) and 1.70 (95% confidence interval bounds: 1.69 - 1.71; R^2^ = 0.9998) respectively. This corresponds to directionally-biased movement at short timescales, although the increasing slope transitioning to smaller scaling exponent of 0.30 (95% confidence interval bounds: 0.28 - 0.32; R^2^ = 0.6063) of supergiants illustrates the transition from directed movement at sub-second timescales to confined movement over seconds to tens of seconds timescales (all fits computed using the Curve Fitter application in Matlab). Taken together, the properties of supergiant trajectories, including higher speeds (Fig. 2C) and constrained directed motion (Fig. 2D,E), are consistent with local search patterns, where cells efficiently scout their immediate surroundings, increasing their chance of prey encounter. These putative search patterns are reminiscent of those exhibited by some animals and used in some robotics contexts^30–35^. In other words, *Euplotes gigatrox* shift to a hunting behaviour pattern as they change their cell size and shape accordingly.

**Fig 2.**
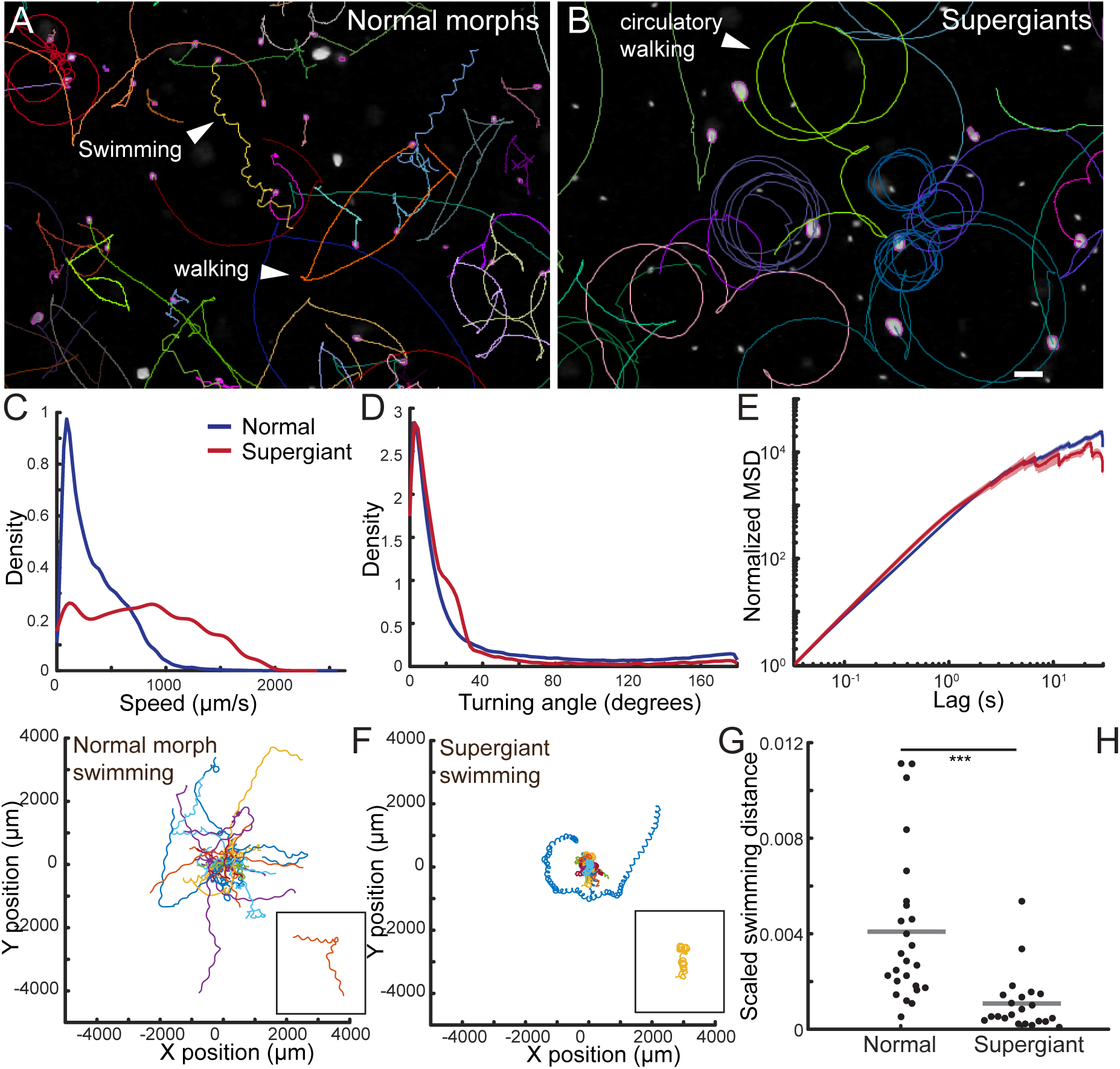
Supergiant formation involves a tradeoff between raptorial feeding and swimming. (A) Representative trajectories of freely-behaving normal morphs display helical swimming and linear walking paths. (B) Representative trajectories of freely-behaving supergiants (from the same sample as A) illustrate curling or circulatory paths, which are less common in normal morphs. Different colours in panels A and B indicate trajectories of different cells and are randomly applied. Scale bar for A and B = 200 µm. (C) A probability-density plot of speeds of freely-behaving normal morph and supergiant cells shows that supergiants tend to move at higher speeds. Speed (y-axis) refers to pooled frame to frame instantaneous speeds (see Methods). (D) A probability-density plot of turning angles along cell trajectories of freely-behaving normal morph and supergiant cells shows supergiant trajectories have different characteristic curvature corresponding to spiraling movements displayed in panel B. For each group of cells, the probability-density estimates in panels C and D were obtained using the ksdensity function in Matlab. (E) Normalized mean square displacement (MSD) of freely-behaving normal morph and supergiant cells shows that each morphotype has similar dispersal characteristics over time, although more rapid decrease in slope of the supergiant MSD curve compared to normal morphs indicates confinement of trajectories over seconds to tens of seconds timescales. Solid lines represent mean MSDs and shaded regions (barely visible) indicate bootstrapped 95% confidence intervals. Measurements for C-E are pooled from 316 normal morph and 42 supergiant cells from three separate samples. (F) Trajectories of swimming normal morphs after they have been gently displaced from the coverslip on which they had been walking display expected extended helical trajectories with occasional turns. These trajectories are pooled from 25 different cells from three different samples and are mean centered. (G) Trajectories of swimming supergiants after they have been gently displaced from the coverslip they had been walking on show compact helical swimming and tumbling. These trajectories are pooled from 22 different cells from three different samples and are mean centered. Colours in F and G are applied randomly and designate different individual cells. (H) Scaled swimming distances calculated by taking the ratio of the distance from the start to the end of the trajectory divided by the total integrated distance traveled by normal morphs and supergiants displayed in panels F and G. Gray bar indicates the mean. *** indicates p<0.001 by two-sample t-test.

We hypothesized that the larger bulk, more irregularly shaped cell bodies, and denser composition all reduce the efficiency of swimming in supergiants. To test this, we gently disturbed cells that were walking on glass-bottomed Petri dishes by pipette aspiration or by shaking the dish. We then recorded the resultant trajectories of cells as they moved through the fluid (Video S5). Supergiants do not appear to be able to swim very far, often tumbling while moving in an undirected fashion, in contrast to the directed, helical swimming of normal morphs documented in this and other *Euplotes* species^28,36^ (Fig. 2F,G, Video S5). Scaled swimming distances, calculated by taking the ratio of the integrated distance traveled by the cell to the total distance traveled from the start to the end of the swimming trajectory, are significantly smaller for supergiants compared to normal morphs (Fig. 2H). These results suggest supergiants have reduced swimming performance in terms of long-range dispersal ability. In principle, swimming allows cells to better explore 3D environments, whereas walking is restricted to surfaces. Supergiant formation therefore may represent a tradeoff where cells can feed on locally available prey items that would otherwise be inaccessible while sacrificing motility over extended distances, perhaps as a strategy to capitalize on ephemeral local conditions. This adaptation is a departure from the more commonly reported “swarmers” or dispersal forms of ciliates and other protists, where under stressful conditions cells become more efficient long-distance swimmers, supposedly for rapid dispersal to favorable environments^37–39^.

### Supergiants result from differentially regulated and reversible developmental pathways

We next sought to investigate conditions under which supergiants form. Many organisms, including protists, undergo life history transitions in response to environmental cues. Indeed, phenotypic plasticity in response to predators, including identification of specific signals in the form of predator-secreted compounds, has been described in several species of *Euplotes*^40–42^. Cannibal formation triggered by environmental conditions has also been reported in other ciliates, notably *Blepharisma*^20^. Starvation is generally proposed as the inducing trigger, although systematic studies have rarely been undertaken. While we observed a consistent yet sporadic appearance of supergiants in our cultures, no obvious trend in culture conditions or timing was apparent. We therefore tested a range of environmental variables including prey abundance, prey type, salinity, temperature, population density, and conditioned media (see Table S1). Contrary to previous reports^22^, these tests were inconclusive, with supergiants continuing to form sporadically under various conditions. To test the dynamics of giant appearance under more controlled conditions, we grew cells in different concentrations of red algae culture medium (RA).

This growth medium serves as a carbon and nutrient source for bacterial prey co-cultured with *Euplotes* as well as a nutrient source for *Euplotes* itself. We found that supergiants consistently began to form as a population entered stationary phase following exponential growth, then increased in abundance before ultimately disappearing (Fig. 3A,B).

**Fig 3.**
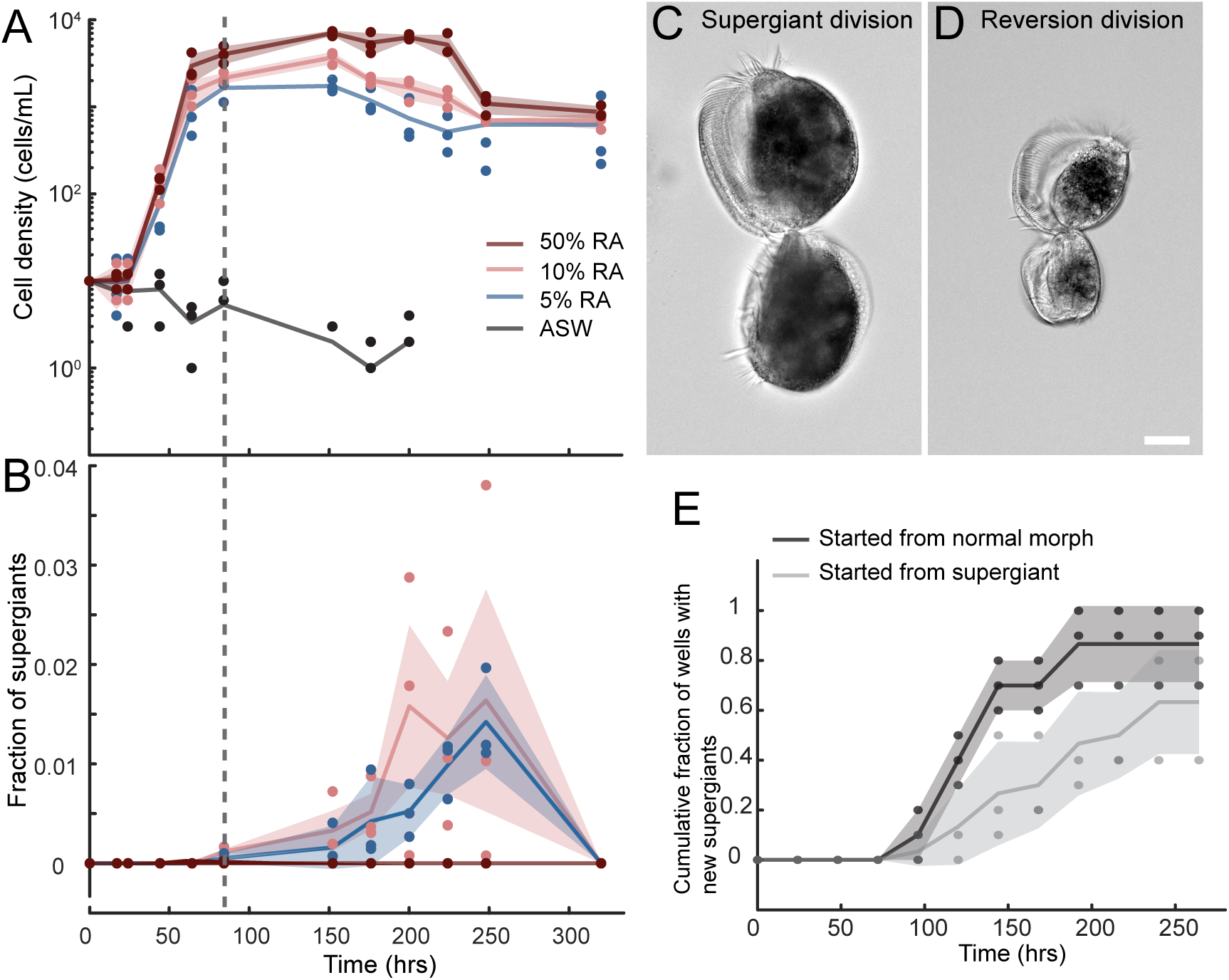
Supergiants form and revert under specific environmental conditions in a history-dependent manner. (A) Growth curves of cells in different media enrichment conditions, denoted by different coloured curves and points. (B) Fraction of supergiants over time corresponding to the same cell populations in panel A. In both A and B, points indicate independent experiments from three trials, solid lines are means, and shaded region indicates standard error of the mean. Note that supergiants only appeared in 5% and 10% red algae media (RA). The vertical dashed line indicates the first appearance of supergiants. (C) Supergiants can propagate supergiants by dividing approximately equally, as depicted in this representative example of cells just prior to separating during cytokinesis. (D) Supergiants revert to normal morphs by a series of asymmetric cell divisions. This image captures an intermediate stage of this reversion process just prior to cells separating during cytokinesis. Scale bar for C and D = 25 µm. (E) Cumulative fraction of wells in a well plate that contain supergiants in populations started from either a single normal morph or a single supergiant in 10% RA media shows that populations started from supergiants form new supergiants more slowly and are less likely to produce supergiants overall. Each experiment involved ten wells seeded by an individual cell; points indicate results from an individual experiment, solid lines represent the mean, and shaded regions represent the standard error of the mean.

Supergiants did not form in the absence of nutrients and cell growth and division (Fig. 3B). Surprisingly, they also did not form under high nutrient conditions (Fig. 3B). Prior to the appearance of supergiants, we often observed cells of intermediate size with extended AZMs and rapid motility (Fig. S3, Video S6). These cells can also feed cannibalistically (Video S6) and may represent a step in the supergiant developmental pathway. In addition to arising from a developmental process from normal morphs (Videos S7), we also found that supergiants can propagate themselves through approximately equal cell divisions (Fig. 3C). Lastly, we documented the process of reversion to the normal cell type, which takes place through asymmetric cell divisions (Fig. 3D). During all these experiments, supergiants never amounted to more than 5% of the population, which suggests that supergiant formation is a low-probability life history transition.

A less explored possibility is that internal cellular states, in addition to external prompts, play an essential role in triggering supergiant formation, as is the case for life history transitions in some other protists^39^. To investigate whether the disposition to form supergiants is history-dependent, we picked individual cells of normal morphs and supergiants and placed them in wells containing either RA medium at a concentration compatible with supergiant formation, or artificial seawater (ASW) free of other nutrient supplements. In both ASW and RA conditions, all supergiants reverted to normal morphs within 24 hours (n = 10, N = 3 for both conditions). In ASW, individual supergiants produced up to nine normal morphs in 24h, and up to 16 normal morphs over 120 hours, whereas individual normal morphs underwent a maximum of one binary cell division in 24h and then stopped dividing. In RA, cells proliferated as expected based on standard culture conditions. Interestingly, supergiants formed earlier and with higher overall frequency in populations started from normal morphs compared to those starting from supergiants (Fig. 3E). These results all support the hypothesis that supergiant formation depends on the internal cellular state, and suggest that supergiant reversion may trigger a latency or cooldown period, likely by temporarily suppressing the pathways that promote the transition to supergiants. Supergiant reversion itself depends on specific environmental stimuli, occurring both in the absence of prey and when small prey is abundant, conditions that also delay the formation of new supergiants (which would likely be maladaptive under these conditions).

Taken together, these results are consistent with supergiant formation being a low-probability life history transition that occurs after a period of recent cell proliferation when small prey items are not too abundant and large prey items are present. Supergiant formation, reversion, and propagation therefore are cyclic regulated processes, controlled by environmental conditions as well as internal cell states.

### Development of *Euplotes gigatrox* is linked to changing gene expression profiles

To investigate the molecular basis of supergiant formation and reversion, we sequenced single cell transcriptomes from 41 *E. gigatrox* cells corresponding to supergiants (n = 9), normal morphs (n = 10), normal morphs that had recently reverted from the supergiant state (n = 10), and ‘intermediates’ (n = 12) whose appearance did not match that of either normal or supergiant cells (Fig. S1), and that we hypothesize represent an heterogeneous group of transitional states. After all sequences were clustered and contamination filtered out, 35,567 distinct transcripts were identified for a total length of 38.3 Mbp, corresponding to 27,557 protein-coding genes (Table S2).

An initial count matrix (Table S3) was used to examine the variance and distribution of genes represented in different libraries through principal component analysis (PCA; Fig 4A). PCAtest^43^ confirmed the overall statistical significance of the analysis as a whole and of the first six principal components (PCs) (Table S4). PC1 and PC2 combined covered 43.6% of the total variance. Normal and supergiant cells both form strongly supported clusters, and more surprisingly so do normal morphs that recently reverted (Fig 4A), demonstrating a molecular basis for the latency period observed during growth experiments. The pattern is supported by a global ANOSIM test on the whole data set (*R* = 0.616, *p* = 0.001), and even more strongly when ‘intermediate’ cells, which are expected to form a less coherent category, were excluded (*R* = 0.8537, *p* = 0.001), as well as by pairwise ANOSIM tests (Table S4). These results suggest that the processes of cellular differentiation to form supergiants, and de-differentiation to reverted and normal stages, are both underpinned at the molecular level by modulating gene expression profiles.

**Fig 4.**
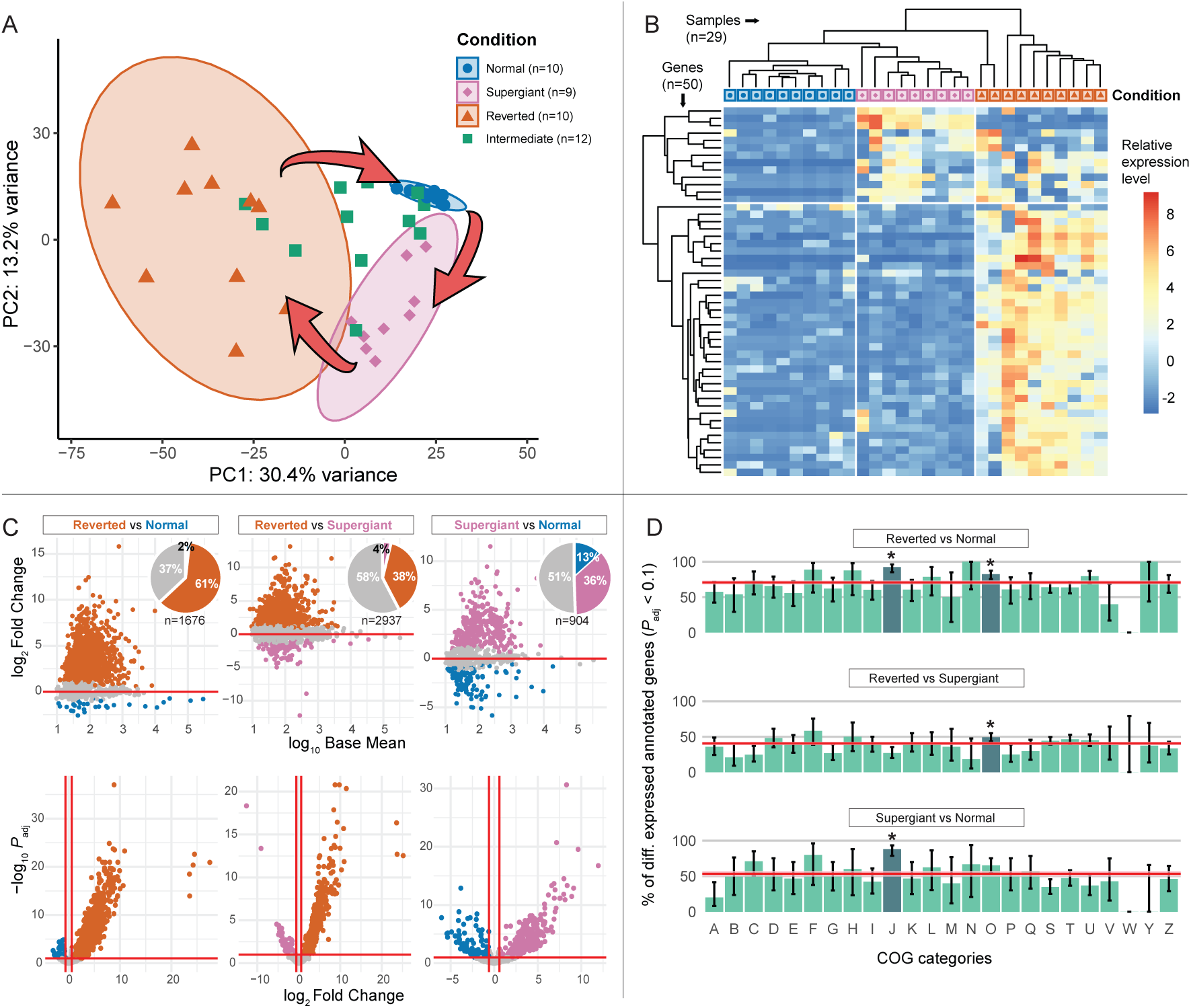
The developmental cycle of *Euplotes gigatrox* is reflected by changes in the expression profiles of two sets of protein-coding genes. (A) Principal component analysis of the top 500 genes in the rlog-transformed count matrix with the highest variance across samples: each dot corresponds to a different single-cell; 95% confidence ellipses highlight three distinct expression profiles corresponding to the three developmental conditions (data on cells displaying intermediate phenotypes is also reported). Arrows are used here to underline the inferred direction of the three-step regulatory cycle of *E. gigatrox*’s development. (B) Hierarchical clustering and heatmap on rlog-transformed count values displaying the 50 genes (rows) whose expression levels have the highest variance across samples (columns). Values represent their deviation from the gene’s average across all samples. (C) Pair-wise differential expression analysis between three developmental conditions; each dot represents one protein-coding gene. MA-plots (top): approximate posterior estimation for generalized linear model shrinkage estimator has been applied to reduce the noise associated with log Fold Changes expected from low count genes^53^. Bottom: volcano plots. Datapoints in gray indicate genes which are not significantly differentially expressed (*P*_adj_ > 0.1 or |log_2_ Fold Change| < 0.6); datapoints in other colours indicate genes which are significantly upregulated in one developmental stage compared to the other. (D) Barplots showing the result of functional enrichment analyses: proportion of differentially expressed genes between each pair-wise comparison, which could be annotated within at least one cluster of orthologous group (COG) functional category (following [45]). Proportions of differentially expressed genes (*P*_adj_ < 0.1) calculated across all COG categories are represented by red lines. Genes involved in translation, ribosomal structure and biogenesis (J), and post-translational modification, protein turnover, and chaperones (O) are shown here as significantly enriched under certain conditions. *P*_adj_= adjusted *p*-value.

A gene-cluster analysis (Fig. 4B) of the 50 genes with the highest expression variance associated with the three distinct developmental conditions (normal, supergiant, and reverted) identified two sets of differentially expressed genes: one set is switched on during the differentiation to supergiant, and partially lingers in the reverted stage, while a second set is upregulated in reverted cells only, and it likely accounts for the latency period. The same pattern is retrieved, albeit less distinctly, when the 500 genes with the highest variance were analysed, except that a higher fraction of genes appeared to be slightly upregulated in supergiants relative to normal cells (Fig. S4).

The putative functions of these genes were assessed from 7,900 out of 27,557 predicted coding sequences annotated using the eggNOG v5 database and 18,871 coding sequences annotated with InterProScan (Table S5). To detect genes that are most relevant for the transitions, we first identified those that contribute more to the variance in the first two components of the PCA analysis by measuring their loading score indices and correlation significance (Table S6). Twenty genes with the highest positive and 20 with the lowest negative loadings scores were selected for both PC1 and PC2, and, for each gene, their expression levels were compared across the different conditions (Table S7).

Genes involved in chromatin structure and dynamics were the most represented category, including genes coding for histones H2A, H2B, H4, and a protective telomere end-binding protein, all of which were upregulated in supergiant and reverted cells compared to normal cells. The annotated gene with the absolute highest loading score codes for a cyclin-dependent kinase regulatory subunit, which also appeared downregulated in normal cells. These observations suggest strong regulatory ties between cell differentiation and genes commonly associated with the cell cycle. Interestingly, a protein displaying a homeodomain, typical of transcription factors involved in animal and plant development^44^, is also overexpressed in supergiant and reverted cells. Among the genes highly expressed in supergiants only, there were genes coding for cytochrome P450, an ABC transporter, and a protein regulating the pyridoxal 5’ phosphate intracellular homeostasis. Genes distinguishing reverted cells were mostly downregulated in the other conditions and included an NADP-dependent oxidoreductase potentially involved in post-translational modification or protein turnover, a NAD-dependent epimerase/dehydratase, translation initiation factor IF2B/IF5, a lariat debranching enzyme, and an adenosine/adenine deaminase.

The total number and extent of differentially expressed genes between each developmental condition pair was visualized though pairwise matrices, each including a different sub-selection of genes (Table S8). No less than 42% of all genes are differentially expressed in each comparison, and the differences are both wide and highly statistically significant (Fig. 4C), suggesting that many genes are involved in the transitions. Noticeably, in reverted cells relative to normal cells, most genes were upregulated, reflecting the same pattern as the cluster analysis (Fig. 4B).

To evaluate whether functional clusters of orthologous genes (COG) categories^45^ were especially enriched in genes that are differentially expressed between developmental conditions, we compared the proportion of such genes across COG categories with that of the background for each condition pairing (Table S9). At least one of two COG categories always appeared to be enriched in differentially expressed (and annotated) genes (Fig. 4D): genes involved in translation, ribosomal structure and biogenesis (J), and genes involved in post-translational modification, protein turnover, and chaperones (O). Specifically, the J category is significantly enriched in genes differentially expressed when either supergiant (88.0% Wilson score CI: 78.7%–93.6%, n = 75) or reverted cells (92.4% Wilson score CI: 85.1%–96.3%, n = 92) are compared to normal cells (two-tailed *Z*-test, *p* < 0.00001). In contrast, the O category is significantly enriched in differentially expressed genes when reverted cells are compared to either supergiant (49.3% Wilson score CI: 43.5%–55.1%, n = 280; two-tailed *Z*-test, *p* = 0.006) or normal cells (82.3% Wilson score CI: 75.6%–87.4%, n = 158; two-tailed *Z*-test, *p* = 0.002). Other COG categories showed no significant difference from the baseline.

## Conclusions

*Euplotes gigatrox* is a newly described ciliate that regulates the reversible development of supergiant cannibals in response to physiological and environmental cues. Supergiants are morphologically, behaviourally, and transcriptionally distinct from normal cells, adopting a novel feeding strategy to exploit larger prey, including genetically identical kin. The process appears to have been fine-tuned during evolution, as shown by details like a cooldown period in recently reverted cells that is underpinned by regulated gene expression, or the coordinated change to a morphology amenable to conspecific-predation at the same time as a change in foraging patterns that turns supergiants into better hunters, but worse swimmers. The regulated yet apparently stochastic trigger of the transition, along with the associated tradeoff between dispersal versus the exploitation of a new trophic niche, suggest that the formation of supergiants could serve as a bet-hedging strategy^46^ for a subset of cells in a recently growing population as it reaches carrying capacity.

Inducible trophic polymorphism through morphological change occurs in diverse eukaryotic lineages including metazoans, amoebozoans, and other ciliates^47–51^. In many cases, it is linked to diminishing resources^25,47,49,50^. Unlike in animals, the morphogenetic processes involved in unicellular protists must take place within a single plasma membrane and continuous cytosol, a topic about which little is known^17,52^. *Euplotes gigatrox* is hence a case study at the interface between ecological and cytological questions, and this work provides a framework on which to build mechanistic studies to address both. By taking an integrative approach combining morphometric, behavioural, experimental, and molecular analysis, this framework can also be a foundation for a more general understanding of developmental processes in unicellular organisms, including their behavioural, ecological, and evolutionary implications. There are certainly many fascinating open questions, even in this system. What specific cues induce supergiant induction and reversion? The transitions are regulated and linked to environmental conditions, but how cells actually sense abundance and prey type, or what specific external or internal signals respond to environmental conditions are all major unanswered questions. The current data suggest direct encounters with prey items play a role, but how an apparently stochastic developmental switch is regulated and how it is maintained over ecological and evolutionary timescales remain open questions. Similarly, raptorial giant cells have evolved multiple times in *Euplotes* and other lineages, raising questions about how they originate in natural contexts and under which conditions selection favours them over other strategies associated to low resource availability; re-isolation and direct observation of cells in the field could help to address these questions. We have identified clusters of genes that are associated with specific stages, but the functional pathways remain untested, so how the modification of specific processes might play a role in the evolutionary diversification of cellular form and function is yet to be determined.

Although phenomena that are controlled by developmental mechanisms similar to those regulating supergiant formation are likely widespread among eukaryotes, their mechanistic bases, functional roles, and natural environmental contexts all must be better understood to clarify their ecological and evolutionary significance. *Euplotes gigatrox* serves as a vivid illustration of unexplored diversity among protists and the potential this holds for shedding new light into fundamental biological principles.

## Methods

### Initial isolation and sustained culture of *Euplotes gigatrox*

Cells were first collected by scraping filters for the water table system at CARMABI research station on the Caribbean island of Curaçao, which draws in seawater from a near-shore environment. Filter scrapings were deposited into vented cap T25 culture flasks (Corning, Corning, NY, USA) and initially kept at room temperature for two weeks before *Euplotes* cells were first directly observed. Cells were picked along with co-occurring bacteria and protists and placed into T25 culture flasks containing 10 mL of 10% Cereal Grass Media (CGM3; as in [54]) kept in a VWR Scientific model 2005 low temperature incubator at 22 °C. The ciliates were observed to proliferate under these conditions, and individual cells were then picked into T25 culture flasks with 10% CGM3 along with co-isolated bacteria to establish strains E1 and E2, and incubated at 22 °C. Cell growth was monitored and after approximately three weeks E1 and E2 were passaged at a 1:10 dilution into 10 mL of 25% CGM3 in T25 culture flasks and kept at 22 °C. Cultures were maintained under these conditions, with visual inspection and passaging approximately every two to three weeks. Supergiants were observed sporadically during this time. While E2 played a role in initial observations, a co-isolated amoeba contributed to less reliable *Euplotes* growth by also feeding on prey bacteria, and therefore strain E1 was used for subsequent experimental work.

For sustained, long-term culture of strain E1, cells were transferred to 15% red algae stock media (RA; as in [55]). Cultures were maintained in T25 flasks at ambient room temperature conditions covered loosely by plastic wrap to reduce evaporation. Cells were passaged approximately every two weeks at a 2:10 dilution into 10 mL fresh media. Under these culture conditions, supergiants were reliably observed.

### Light microscopy and morphometric analysis

Cells for morphometric analysis were imaged in Fluorodishes (World Precision Instruments FD35-100, Sarasota, FL, USA) by differential interference contrast microscopy using a 20x Plan Apochromat Lambda D (0.8 NA) Nikon objective on a Nikon ECLIPSE Ti2 inverted microscope (Nikon, Shinagawa, Tokyo, Japan) with a Photometrix Kinetix camera (Teledyne, Waterloo, ON, Canada). Cell measurements were performed manually in Fiji^56^.

For routine observation and documentation (e.g. Fig. 1B,L), cells were imaged in T25 culture flasks by brightfield or phase contrast using a 20x Plan Fluor ELWD (0.45 NA) Nikon objective on a Nikon ECLIPSE Ts2 inverted microscope with a Nikon Z6 camera.

### Scanning electron microscopy

Cells were deposited onto plasma-cleaned coverslips (#1.5, 12 mm, Electron Microscopy Sciences, 72290-04, Hatfield, PA, USA) coated with poly-D-lysine (Sigma Aldrich P6407-5MG, Burlington, MA, USA) and fixed for two hours at 4 °C in 4% paraformaldehyde and 2.5% glutaraldehyde in PHEM buffer and artificial seawater (ASW, ASTM D 1141, Ricca Chemical Company, 8363-5, Arlington, TX, USA).

Samples were then rinsed with fresh water and then dehydrated by ethanol dilution series according to the following steps: 25% ETOH in water (5 min), 50% ETOH (5 min), 75% ETOH (5 min followed by overnight incubation at 4 °C), 95% ETOH, 100% ETOH (10 min, repeated for a total of three times). Samples were critical point dried for 80 minutes using a Leica EM CPD300 (Leica Microsystems, Wetzlar, Germany) under a slow automated setting and sputter coated with 6 nm of iridium using a Leica EM ACE600 before imaging using a Phenom Pharos G2 desktop field emission gun scanning electron microscope (Thermo Scientific, Waltham, MA, USA).

### Phylogenetic analysis

The full-length small subunit (SSU) rRNA sequence was extracted from the reference transcriptome (see below), and confirmed to be identical to a sequence of the SSU rRNA gene obtained through DNA extraction and PCR (data not shown). It was aligned with 77 representative orthologs from other *Euplotes* species collected from GenBank as well as 11 sequences from related genera (to serve as outgroup) using the linsi option in mafft^57^ v7.520. The alignment was trimmed at both ends to remove sites with more than 50% missing data, resulting in a 89 taxa by 2,168 sites character matrix. Maximum likelihood phylogenetic inference was performed on IQ-TREE^58^ v2.2.2.7, using the GTR+F+I+G4 model as suggested by all information criteria, and running 1,000 nonparametric bootstrap pseudoreplicates for support.

### Cell motility analysis

For motility analysis, 2 mL of cell culture were taken directly from culture flasks and placed in fluorodishes. This amount of fluid ensured sufficient depth to allow unconstrained swimming behaviour. Cells were imaged in dark field on a Zeiss Axio Zoom.V16 microscope with a Zeiss 1.0x Plan Neo Fluor objective (NA 0.25) and a Zeiss AxioCam 305 colour camera (Zeiss Microscopy, Oberkochen, Germany). Videos were acquired at 60 frames/s and were pre-processed in Fiji by first subtracting the individual average pixel value over the entire video (background) from each video frame and then smoothed by applying a median filter with a 4 pixel radius. Tracking was then performed with the TrackMate Fiji plugin^59,60^ using the following settings: threshold detector for blob detection, LAP tracker frame to frame linking = 10 px, gap closing = 10 px, and frame gap = 10 frames. To remove mis-identified tracks, a post-processing filter for tracks less than 40 frames and speeds greater than 16 px/frame was applied. Supergiants were discriminated from normal morphs based on size by applying a post-processing filter with a threshold radius of 7 pixels, where blobs above this size were designated supergiants and those below were designated normal morphs. Visual inspection confirmed that this threshold never led to the misidentification of supergiants.

Trajectories were then smoothed using a Savitzky-Golay filter with polynomial order 2 and frame length 5 implemented using the sgolay function in Matlab (R2023b, Mathworks, Natick, MA, USA). Instantaneous speeds were calculated at discrete time *t* as 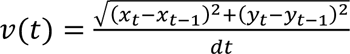 where (*x_i_*, *y_i_*) are the coordinates at frame *i* and *dt* is the interframe time interval. Turning angles along trajectories were calculated by θ = *atan2(|<u>p</u> × <u>q</u>|, <u>p</u> • <u>q</u>*) where *p* = *r*_t_ – *r*_t-5_ and *q* = *r*_t+5_ – <u>*r*</u>_t_ are vectors approximately tangent to adjacent positions along the trajectory where *r*_n_ denotes a cell position at frame *n*, following [15]. Mean square displacement was calculated by 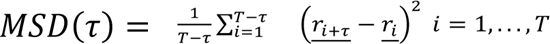 for time lag τ, incremented by the frame rate, where *T* is the time for the final frame of the trajectory. MSDs were normalized by dividing by the value at the first point of the MSD curve. 95% confidence intervals were determined by bootstrapping using the bootci function in Matlab with 1000 samples.

For swimming comparison experiments, cells were gently displaced from the coverslip surface of fluorodishes by gentle shaking of the dish or pipette aspiration while simultaneously recording on the Axio Zoom microscope (setup detailed above). Tracking was performed in trackmate as described above, and each trajectory of a swimming cell was manually evaluated to determine the beginning and end of unconstrained swimming, free of apparent influence by fluid currents introduced by knocking cells from the coverslip.

### Supergiant induction experiments and analysis

All supergiant induction experiments presented in the main text (Fig. 3A,B,E) were conducted in 24-well plates (Fisher Scientific 09-761-146) in 2 mL growth media or ASW. For experiments reported in Fig. 3A, 10 normal morph cells were picked from a culture one week after a fresh passage, placed into each well, and allowed to proliferate at room temperature. Plates were sealed with two layers of parafilm to reduce evaporation. Cell counts were performed manually, by dividing wells into quadrants for normal morphs, and directly for supergiants; the average of three counts was taken. Cell counts were performed using a Zeiss Stemi 508 Microscope under darkfield illumination. For experiments reported in Fig. 3B, individual normal morph or supergiant cells were picked into individual wells, and plates were monitored every day for the presence of supergiants. See Table S1 for methodological information on additional supergiant induction experiments not reported in main text figures.

### Single-cell collection for transcriptomics

Supergiant cells were transferred with a pulled glass micropipette under a Zeiss Stemi 508 Microscope into separate wells of a glass spot plate with depressions, in autoclaved filtered seawater and with no food source. Cell division and consequent reversion to normal morphs were monitored by checking the wells every 2 hours four times. After ∼24 hours, when all cells had reverted to the normal phenotype, 10 of them were individually collected, washed in sterile medium to further reduce contamination, and placed each into a 0.2 mL tube containing 2 μL of lysis buffer^61,62^. Additionally, 10 normal and 9 supergiant cells were individually collected from the original cultures and also placed in lysis buffer. Finally, 12 cells with intermediate or ambiguous features were also collected and isolated, 6 from each of two separate cultures of the same clonal strain. The 41 collected cells were stored at −70 °C until further use.

### RNA-seq, reference transcriptome co-assembly, and contamination removal

First, frozen samples were subjected to at least four freeze-thaw cycles to facilitate cell lysis^63^. Then, cDNA was extracted from each sample and purified following the Smart-seq2 protocol^62^. The obtained cDNA yield was quantified with Qubit^TM^ dsDNA HS Assay Kit and fluorometer (Thermo Fisher Scientific). The cDNA samples were submitted to the Sequencing + Bioinformatics Consortium of the University of British Columbia (BC, Canada). There, length composition and quality of the samples were assessed with an Agilent Bioanalyzer (High Sensitivity DNA Assay); libraries were prepared for sequencing using Illumina DNA Prep; paired-end sequencing was carried out on a NextSeq 500 platform (2×150 bp).

Prior to assembly, all reads were pre-processed with Trimmomatic^64^ v0.39 to remove adapters and for quality filtering. Then, rnaSPAdes^65^ v3.15.5 was used to co-assemble reads from all samples into one preliminary transcriptome, as an initial step towards constructing a comprehensive reference. An initial set of 57,651 transcripts (each longer than 300 bp) and total length of 56.8 Mbp was obtained, from which coding regions were predicted through TransDecoder v5.5.0 with homology searches^66^; the ‘Euplotid’ genetic code was selected, which differs from the universal genetic code as it replaces the UGA codon’s stop signal with a cytosine^67^. The predicted coding sequences were clustered with cd-hit-est^68^ setting a lower threshold of 90% identity, to reduce redundancy and limit the chance of multiple mapping during the alignment of the reads to the reference transcriptome. The aminoacidic sequences were blasted with BLASTp to identify the likely source of each transcript and limit any potential contamination: all transcripts hitting Alveolata were kept; other sequences with less than 80% identity were kept if their GC content ranged within the 10^th^ and the 90^th^ percentile of the Alveolata distribution (36% and 42% respectively). All hits to Metazoa, Fungi, Stramenopila, Viridiplantae, *Pseudomonadota*, *Terrabacteria*, *Bacteroidota*, *Planctomycetota*, and viruses were filtered out. After sequences were clustered and potential sources of contamination were removed, the final reference transcriptome retained 35,567 transcripts and had a total length of 38.3 Mbp, corresponding to 27,557 protein-coding genes. Busco^69^ v5.4.3 and its ‘alveolata_odb10’ database (n = 171) were used to estimate and compare the completeness and the level of duplication of the initial and final reference transcriptome. While the BUSCO completeness score expectedly decreased from 71.3% to 57.9% throughout the filtering process, the proportion of single-copy markers increased from 45.6% to 56.1%, and that of duplicated markers plummeted from 25.7% to 1.8%, thus improving univocal read alignment to the reference. The proportions of fragmented matches remained roughly unchanged (18.7% and 19.3% respectively).

### Transcriptomics - data analyses in R

All analyses were conducted on R v4.2.1 and v4.3.2. Transcript quantification into count matrices, gene-level exploratory analysis and visualization, and differential expression analyses were carried out using the Bioconductor^70^ v3 R packages and guidelines described by [71], with few adjustments. Briefly, reads from each sample were separately aligned to the reference transcriptome using bowtie^72^ v2.5.3; gene models were defined from the GFF3 file obtained from the TransDecoder coding region prediction described earlier; sample information was added to a *SummarizedExperiment* (SE) object, choosing a developmental condition as reference level; the SE object was then converted into a *DESeqDataSet* (dds) object (count matrix) for further analysis. The dataset was then pre-filtered keeping only rows that have a count of at least 10 for a minimal number of samples. The regularized-logarithm (rlog) transformation was applied to make the data homoscedastic for successive exploratory analyses.

### Transcriptomics - exploratory analyses

From the rlog-transformed count matrix, the 500 genes with the highest variance across samples were selected for a principal component analysis (PCA), performed through the built-in R function *prcomp()*. The *PCAtest* package^43^ v0.0.1 was used to test the PCA overall significance, that of each principal component (PC), and that of the correlation of gene variables to the PCs based on 1,000 bootstrap replicates and 1,000 random permutations. The minimum number of significant PCs to retain was further assessed through Horn’s parallel analysis^73^ implemented within the *paran* v1.5.3 package^74^ using 10,000 iterations. Among the six retained PCs, the first two combined covered a large fraction of the total variance (43.6%), so they were selected to plot their score values using ggplot2 v3.5.0 from the tidyverse R collection^75^. The clustering patterns of the data points and their significance were evaluated with an analysis of similarity (ANOSIM) test implemented with the *vegan* v2.6-4 package^76^, setting 999 unrestricted permutations: six tests were performed for each pair-wise comparison between the examined groups (normal, supergiant, reverted, intermediate); for each, a distance matrix was calculated based on Euclidean distances between the data points; *R* statistics and their respective *p-* values were then obtained for each pair. To correct for the increase of the type I error rate due to multiple testing, *p-*values were adjusted using the Bonferroni correction. To measure the clustering tendency of data points as a whole, the ANOSIM test was also performed across all groups.

The top 50 and the top 500 genes of the rlog-transformed count matrix with the highest variance across samples were selected for hierarchical clustering to identify patterns of expression (if present) between the different developmental conditions. Data from the intermediate-looking cells were disregarded from this and further analyses, because of their lack of unambiguous characterization. The clustering was carried out using pheatmap^77^ with the default clustering method (i.e. complete linkage).

### Pair-wise differential gene expression analyses

Differences in the expression levels between developmental conditions of *Euplotes* were examined individually by pair-wise comparison through DESeq2^78^ v1.36.0. Precautions were adopted in dealing with potential zero-inflation effects expected in single-cell datasets^79^. For this, the DESeq2 manual recommendations^80^ for single-cell analysis were followed: size factors were calculated using the *scran* package; the Wald test was replaced with the likelihood ratio test for significance testing; other *DESeq()* arguments were adjusted as: useT = TRUE, minmu = 1e-6, and minReplicatesForReplace = Inf.

### Functional annotation and enrichment analyses

Genes predicted from the reference transcriptome were annotated using eggNOG-mapper^81^ v2.1.0-1 with the eggNOG database^82^ v5.0.2. The search step of eggNOG-mapper relies on DIAMOND^83^ v2.0.15. Domain-scale annotations were obtained with InterProScan^84,85^ v5.75 by running all the default analyses. The eggNOG-mapper annotation output for each gene was matched with the results obtained from the differential expression analyses; then, proportions of differentially expressed genes with an adjusted *p-*value < 0.1 were calculated for each cluster of orthologous group (COG) functional category and across all categories. Confidence intervals for each proportion were obtained using the Wilson score interval^86^.

## Acknowledgments

We would like to thank David Booth for the use of his light microscopes and Dyche Mullins for the use of his electron microscope, sputter coater, and critical point dryer. We would also like to thank Kristen Marhaver for giving us access to the filters from which *E. gigatrox* was isolated. Thanks to Vojtěch Žárský for providing initial guidance and technical support in processing transcriptomic data for the analysis of gene differential expression. BTL would like to acknowledge funding from the Merck Fellowship of the Jane Coffin Childs Memorial Fund for Biomedical Research. BTL also thanks Nicole King and Wallace Marshall for tolerating the existence of the initial project that eventually led to this publication and for harboring cell cultures in their labs. PJK acknowledges funding from the Gordon and Betty Moore Foundation (https://doi.org/10.37807/GBMF9201). DG thanks the University of British Columbia for funding this research through the Four Year Doctoral Fellowship Award.

## Supplementary Information

**Fig S1** Example of a *Euplotes gigatrox* cell with intermediate features between the normal and supergiant morphs. Scale bar = 20 µm.

**Text S1 Establishment of *Euplotes gigatrox* sp. nov. and comparison with other *Euplotes* species.** Full rationale for the assignment of the investigated strain to a novel species; species formal establishment and diagnosis.

**Fig S2 Phylogenetic position of *Euplotes gigatrox*.** SSU rRNA gene-based tree of *Euplotes* species, with 11 sequences from other euplotids used as outgroup. *Euplotes gigatrox* belongs to clade A and is closely related to a strain mistakenly identified as *Euplotes parkei* and another assigned to *Euplotes charon*. Numbers associated with nodes represent nonparametric bootstrap support (values below 75% are omitted).

**Video S1 Example of a cannibalistic event.** A supergiant morph captures a genetically identical kin from the same culture. Video plays in real time.

**Video S2 Example of a cannibalistic event including internalization of prey.** Note the cytoplasmic flow starting beneath the oral cavity that accompanies the internalization of the prey cell; the predator cell remains stationary during prey consumption. Snapshots from this video are displayed in Fig. L-L’’’’. Video plays in real time.

**Video S3 Trajectories of freely behaving normal morphs.** Different colours designate individual cells and are randomly applied. Tracking was done using the TrackMate Fiji plugin (see Methods). Video plays in real time.

**Video S4 Trajectories of freely behaving supergiants.** Different colours designate individual cells and are randomly applied. Tracking was done using the TrackMate Fiji plugin (see Methods). Video plays in real time.

**Video S5 Swimming behavior of normal morphs and supergiants that have been knocked from substrate.** Note that many cells remain in contact with the substrate over the course of this experiment. This video is representative of those used to quantify swimming performance and summarized in Fig. 2, where only trajectories of cells swimming after being knocked from the substrate were analyzed.

**Table S1 Summary of inconclusive supergiant induction experiments.** Experiments are grouped in the table by condition tested. All experiments were conducted in glass well plates and 400 µL total volume, except the conditioned media tests, which were conducted in 24-well plates, and the 9.6 cm^2^ surface tests, which were conducted in 6-well plates, both of which contained 2 mL of cell culture. Each experiment was monitored daily over up to 24 days for the appearance of supergiants, which were counted manually. All experiments were conducted in 5 replicates, except temperature experiments, which were conducted in triplicate. For all experiments except for prey, conditioned media, and local density experiments, cells were fed adding 10 µL of a culture of the green alga *Dunaliella* grown to saturation and diluted 1:100 in autoclaved seawater. For prey experiments, cells were fed as specified and the prey organisms, *Duniella* and the bacterium *Klebsiella (Enterobacter) aerogenes*, were diluted in autoclaved seawater. Cultures were grown starting from 100 initial cells in 15% RA medium for the conditioned media and local density experiments, or starting from 100 µL of sterile filtered ASW or media from cultures containing supergiants or not containing supergiants for the conditioned media experiments. For population density experiments, cells were culled three times per week to maintain the stated densities, except for the saturation condition, under which the initial 100 cells were allowed to grow without culling. Together, these experiments suggested that cell proliferation was tied to supergiant formation and that supergiants can form under a range of environmental conditions, although not in a strictly deterministic fashion.

**Fig S3** Morphogenetic progression from an intermediate morph to a supergiant, including a cannibalistic predation event by the intermediate. (A) The first appearance of the intermediate morph in the video. (B) The intermediate captures a prey cell. (C) The intermediate stops moving as it begins to engulf the prey. (D) The prey begins to be internalized, and the intermediate continues to remain stationary. (E) The prey has almost been entirely engulfed. (F) The intermediate moves again after having ingested its prey. (G) The intermediate has now become a supergiant. The cannibalistic event depicted by snapshots here is displayed in timelapse in Video S6. Note that images have been registered to keep the cell in the center of the field of view. Timestamps are hours:minutes:seconds. Scale bar = 50 µm.

**Video S6** Cannibalistic predation by an intermediate morph. Timelapse recording plays at 10x real time.

**Video S7** Documentation of the appearance of a supergiant. The population consisted initially of only normal morphs. Video plays at 250x real time.

**Table S2** Abundance and length statistics of raw reads. Reads were obtained from paired-end Illumina sequencing of 41 cDNA libraries, each corresponding to a single *Euplotes gigatrox* cell falling in one of three recognizable developmental stages (normal, supergiant, recently reverted), or categorized as an intermediate.

**Table S3** Raw (un-normalized), normalized, and regularized log-transformed count matrices. Each row corresponds to a different protein-coding gene; columns represent distinct single-cell transcriptomes, sorted by developmental category. Raw values indicate the number of read fragments that could be assigned to a certain gene in a certain transcriptome.

**Table S4** Test results. Tests of significance, component retention, and ANOSIM performed on the PCA dataframe based on regularized log-transformed counts. Euclidean distances were calculated for ANOSIM using PCA scores from the first two principal components.

**Fig S4** Extended hierarchical clustering and heatmap of differentially expressed genes. Heatmap obtained using the top 500 protein-coding genes with highest variance across samples from the rlog-transformed count matrix (rather than the top 50, as shown in Fig. 4B). Rows represent individual protein-coding genes, columns represent different transcriptomes (single cells) assigned to a developmental stage. Values represent the deviation from the average calculated for each row.

**Table S5** eggNOG-mapper and InterProScan annotations of protein-coding genes predicted from the reference (co-assembled) transcriptome.

**Table S6** Loading scores, correlation, and significance of 500 protein-coding genes with respect to the first two PCA principal components.

**Table S7** Annotations and normalized counts of top 20 and bottom 20 most relevant and significant loadings with respect to the first two PCA principal components.

**Table S8** DESeq2 pairwise gene differential expression analyses.

**Table S9** Pairwise COG categories enrichment analysis. Frequencies of functional categories were calculated based on abundances obtained from eggNOG-mapper annotations.

